# A small molecule RIG-I agonist serves to adjuvant broad multifaceted influenza virus vaccine immunity

**DOI:** 10.1101/2022.09.20.508779

**Authors:** Emily A. Hemann, Megan L. Knoll, Courtney R. Wilkins, Caroline Subra, Richard Green, Adolfo García-Sastre, Paul G. Thomas, Lydie Trautmann, Renee Ireton, Yueh-Ming Loo, Michael Gale

## Abstract

Retinoic acid-inducible gene I (RIG-I) is essential for activating host cell innate immunity to regulates the immune response against many RNA viruses. We identified a small molecule compound, KIN1148, that directly binds RIG-I to drive IRF3activation to impart the expression of IRF3-target genes, including specific immunomodulatory cytokines and chemokines. KIN1148 does not lead to ATPase activity or compete with ATP for binding but activates RIG-I to induce antiviral gene expression programs distinct from type I interferon (IFN) treatment. When administered in combination with a vaccine against influenza A virus (IAV), KIN1148 induces both neutralizing antibody and broadly cross-protective IAV-specific T cell responses compared to vaccination alone, which induces comparatively poor responses. This robust KIN1148-adjuvanted immune response protects mice from lethal H1N1 and H5N1 IAV challenge. Importantly, KIN1148 also augments human CD8^+^ T cell activation. Thus, we have identified a small molecule RIG-I agonist that serves as an effective adjuvant in inducing non-canonical RIG-I activation for induction of innate immune programs that enhance adaptive immune protection of antiviral vaccination.

**Summary:** Hemann, et. al. identify a small-molecule RIG-I agonist (KIN1148) that directly binds RIG-I for non-canonical activation and adjuvants pandemic and avian influenza virus vaccination. KIN1148 augments broadly neutralizing antibody and T cell responses in mice and enhances human DC maturation and CD8^+^ T cell activation.

## Introduction

Non-self detection of pathogen-associated molecular patterns (PAMPs) by specialized pattern recognition receptors (PRRs) initiates innate immune signaling cascades that result in tailored antimicrobial responses to limit infection. These responses include the production of cytokines, chemokines and/or type I and II interferons (IFNs) that mediate antiviral defense and regulate adaptive immune responses to clear infection and provide long-term memory protection^1, 2^. Specifically, the PRR family of retinoic acid-inducible gene I (RIG-I)-like receptors (RLRs), RIG-I and melanoma differentiation-associated protein 5 (MDA5), are RNA helicases and members of the RIG-I-like receptor (RLR) family. RIG-I and MDA5 are essential for the initial detection of most RNA viruses where they sense and bind to viral RNA in the cytosol of the host cell^3–5^. Upon engaging its cognate PAMP, short, double-stranded RNA or single-stranded RNA containing poly-uridine motifs marked with 5’-triphosphates, RIG-I is released from its basal, autoinhibited state to initiate a signaling cascade that typically culminates in the induction of IRF3/7- and NFκB-dependent transcriptional programs^6–9^. These transcriptional programs include expression of antiviral genes, proinflammatory cytokines, and IFNs that coordinately function to limit virus infection.

Many microbial PAMPs are effective vaccine adjuvants through their innate immune activation properties. For example, the Toll-like receptor (TLR) agonists CpG oligodeoxynucleotides (TLR9) and monophosphoryl lipid A (MPL) (TLR4) are agonists of IRF3 or NF-kB actions in innate immune programming, and are approved for use in vaccines for hepatitis B virus, herpes zoster, influenza virus, and papillomavirus^10, 11^. These PAMP-derived adjuvants target specific, defined immune pathways to elicit their effects. Developing a broad arsenal of adjuvants that target different and specific arms of innate immunity, to be used independently or in combination, could allow for tailoring of the host immune response to individual vaccines for optimal protection against infection. Such a strategy may greatly enhance the efficacy of many subunit vaccines, like those administered to protect against influenza virus infection, which exhibit lower immunogenicity as compared to live, attenuated or inactivated vaccines. Indeed, several adjuvants are currently being investigated to reduce the amount of vaccine needed to provide protection (dose sparing), to elicit broader protection against viral variants not represented in the vaccine, or to enhance protection of immunocompromised populations^12–15^.

Given the robust immune response induced downstream of RIG-I, synthetic PAMP RNAs that engage RIG-I and potently activate the host antiviral response are under development for therapies to control virus infection^16–21^. These RIG-I-agonistic RNAs potentiate dendritic cell (DC) cross-presentation of antigen to cytotoxic T lymphocytes (CTL) and T follicular helper cell induction, leading to enhanced antibody production^17–20^. When administered in combination with a vaccine, these RNAs improve protection of mice from subsequent influenza virus challenge. Similarly, we predicted that small molecules targeting RIG-I may induce innate immune programs that could adjuvant actions.

We previously conducted a cell-based screen in which we identified innate immune-activating small molecules that conferred either antiviral or adjuvant activities^22–24^. KIN1000 and its medicinal chemistry optimized analog KIN1148 belong to a class of benzobisthiazoles selected for further investigation due to their ability to activate IRF3^24^. In this study, we demonstrate that KIN1148 directly engages RIG-I to activate IRF3-dependent innate immune responses, making it the first small molecule RIG-I agonist to be identified. Biochemical studies show that KIN1148 binds to RIG-I to drive RIG-I self-oligomerization and downstream signaling activation in an RNA- and ATP-independent manner. Transcriptional programs induced by KIN1148 treatment exhibit shared and unique signatures to that induced by other RIG-I agonists including Sendai virus (SeV) infection and hepatitis C Virus (HCV) PAMP RNA transfection^7–9^. KIN1148 adjuvants a split influenza A virus (IAV-SV) vaccine at suboptimal dose to protect mice from lethal challenge with a recombinant highly pathogenic avian H5N1 influenza virus, A/Vietnam/1203/2004 HA and NA combined with A/PR/8/34. When administered with IAV-SV, KIN1148 enhanced both humoral and T cell immune responses to IAV-SV. Importantly, *ex vivo* studies show that KIN1148 promoted dendritic cell (DC) maturation and antigen-dependent activation of human CD8^+^ T cells, suggesting its potential efficacy as an adjuvant in humans. Thus, KIN1148 is a small molecule RIG-I agonist and a novel innate immune-inducing adjuvant.

## Results

### KIN1148 is a small molecule agonist of IRF3 signaling that confers adjuvant activity

The cell-based screen that led to the discovery of KIN1000 (Figure 1a) was intentionally designed for identification of IRF3 activating small molecules^22^; the screen was conducted in Huh7 cells, a human hepatoma cell line in which RLR signaling to IRF3 is functional while signaling through the cytosolic DNA-sensing pathway and the TLRs is defective^25–29^. KIN1148 (Figure 1a) was designed as a medicinal chemistry optimized analog of KIN1000^23, 24^. KIN1148 treatment induces phosphorylation of IRF3 and NFκB p65 (Figure 1b), indicative of innate immune activation.

**Figure 1.**
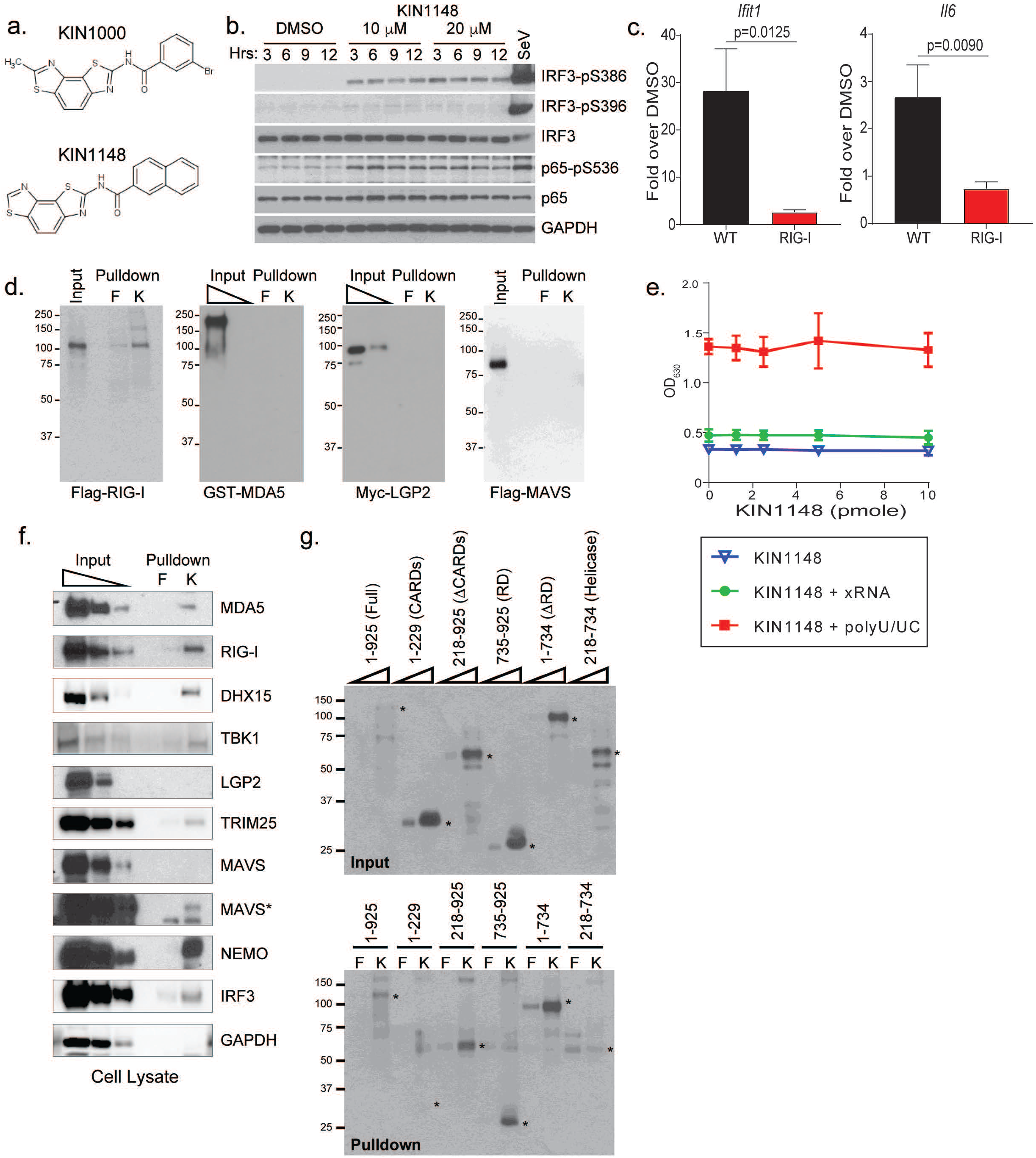
KIN1148 is a small molecule IRF3 and NFκB agonist that binds to and induces RIG-I signaling activation. **(a)** Chemical structure of parent compound KIN1000 and its medicinal chemistry optimized lead KIN1148. **(b)** Western blot analysis of the phosphorylation state of IRF3 and NFκB at time points up to 12 hrs after HEK293 cells were treated with KIN1148 at 10 or 20μM. Included as comparison are cells treated with 0.5% DMSO (vehicle control) and cells infected with SeV at 40 hemagglutinin units (HAU)/mL. **(c)** qRT-PCR analysis of *Ifit1* and *Il6* in WT and RIG-I^−/−^ MEFs at 18hrs after KIN1148 treatment as compared to respective DMSO controls. Fold induction over DMSO control from 3 independent experiments shown. Error bars represent standard deviation. **(d)** Western blot detection of input and pulldown products that associate with either biotin-fluorescein (F) or biotin-KIN1148 (K) beads using recombinant proteins generated using a rabbit reticulocyte lysate *in vitro* transcription and translation system and detected using antibodies specific for the respective epitope tags. **(e)** BIOMOL green colorimetric analysis of free phosphate levels as an indication of RIG-I ATP hydrolysis activity. Purified recombinant RIG-I protein and ATP were incubated with DMSO or increasing concentrations of KIN1148 in the absence and presence of polyU/UC or xRNA. Each point shows the average concentration of free phosphates in each reaction from at least three independent experiments with error bars showing standard deviation. **(f)** Western blot detection of endogenous RIG-I and associating signaling cofactors in the HEK293 whole cell lysate (input) and amongst the pulldown products of biotin-KIN1148 beads (K) or biotin-fluorescein bead alone (F). MAVS* indicates higher exposure blot. **(g)** Western blot detection of various Flag-tagged RIG-I constructs generated by *in vitro* transcription and translation for interaction with biotin-fluorescein or biotin-KIN1148. Asterisks indicate expected location of band of interest across upper and lower blots in **g**.

### KIN1148 is a small molecule RIG-I agonist

To define cellular targets that may engage KIN1148 to mediate innate immune activation, we evaluated innate immune gene expression in mouse embryonic fibroblasts (MEFs) and human alveolar epithelial cells (A549) lacking specific RLRs. Specifically, we measured expression of *Ifit1*, an IRF3-target gene, and *Il6*, a pro-inflammatory cytokine gene encoding IL-6. RIG-I deficient cells demonstrate reduced innate immune gene expression (*Ifit1* and *Il6*) following KIN1148 treatment as compared to wildtype (WT) cells (Figure 1c). KIN1148 induction of innate immune genes is similarly decreased in A549 cells lacking RIG-I as compared to control cells or MDA5 deficient cells (Figure S1a and S1b). Further, we found that biotin-tagged KIN1148 binds to RIG-I in a cell free *in vitro* interaction, but the related RLRs MDA5 or LGP2 do not bind KIN1148 (Figure 1d). Biotin-fluorescein (F) served as a small molecule negative control and did not bind any of the RLR proteins. Additionally, biotin-KIN1148 does not bind directly to Mitochondrial Antiviral Signaling adaptor (MAVS), the essential downstream RLR adaptor protein (Figure 1d). Taken together, these data indicate that KIN1148 directly targets and binds to RIG-I. This interaction is further demonstrated by biotin-KIN1148 capture of endogenous RIG-I from whole cell lysates prepared from HEK293 cells and MEFs (Figure S1c and S1d). Hence, KIN1148 binds to both human and mouse RIG-I to induce innate immune activation.

Upon engaging PAMP RNA, RIG-I hydrolyzes ATP to undergo conformational change and initiate downstream signaling^30–32^. We assessed the ability of KIN1148 to promote RIG-I hydrolysis of ATP by incubating purified recombinant RIG-I with DMSO or increasing concentrations of KIN1148 in the presence of ATP. KIN1148 alone does not lead to hydrolysis of ATP (Figure 1e). Interestingly, KIN1148 does not alter RIG-I ATPase activity when added in increasing amounts to either polyU/UC (a validated RIG-I PAMP RNA derived from the hepatitis C virus (HCV) genome^8^) or xRNA (an RNA of equivalent length from a region within the HCV genome adjacent to polyU/UC that does not activate RIG-I signaling) (Figure 1e). Moreover, while KIN1148 competed with itself for biotin-KIN1148 capture of purified recombinant RIG-I, the addition of ATP or AMP-PNP did not compete for biotin-KIN1148 capture of RIG-I (Figure S1e).

The conformational change in RIG-I leads to the release of its tandem N-terminus Caspase Activation and Recruitment Domains (CARDs) from autoinhibition allows RIG-I to form homo-oligomers. RIPLET binds to oligomerized RIG-I leading to ubiquitination and RIG-I translocation to intracellular membranes to interact with MAVS^33^. This in turn triggers MAVS filamentation and recruitment of signaling cofactors to form an innate immune signalosome on MAVS that leads to innate immune activation^34–40^. To define KIN1148 activation of RIG-I, biotin-KIN1148 pulldown assays were conducted using buffers of lower stringency to recover RIG-I plus associated factors from whole cell lysates. Under such conditions, we recovered MAVS in addition to RIG-I from whole cell lysates (Figure S1c). We additionally identified many of the cofactors that comprise the RIG-I signalosome and which are required for RIG-I signaling to the innate immune response^3, 5, 34, 35^, including DHX15, TBK1, TRIM25, NEMO or IKKγ and IRF3 (Figure 1f).

To determine where KIN1148 may bind to RIG-I, we investigated binding of biotin-KIN1148 with several RIG-I deletion mutants. Biotin-KIN1148 captured all RIG-I constructs that contain either the C-terminus repressor domain (RD) or the conserved DEAH helicase motifs, but not the construct that consists of the CARDs alone (Figure 1g). The data suggests binding sites for KIN1148 are within the RD and the helicase domains of RIG-I but not within the CARDs.

### KIN1148 confers an innate immunity gene expression profile consistent with RIG-I activation

We next aimed to delineate the gene expression profiles of human macrophage-like THP-1 cells treated with KIN1148 or K1000 compared to cells treated with known innate immune agonists, including the TLR4 PAMP lipopolysaccharide (LPS) and IFNβ for Janus Kinase (JAK)/Signal Transducers and Activators of Transcription (STAT) mediated signaling downstream of the IFNα/β receptor (IFNAR). SeV-infected cells and polyU/UC RNA transfection served as controls for RIG-I- and MAVS-specific responses (Figure 2).

**Figure 2.**
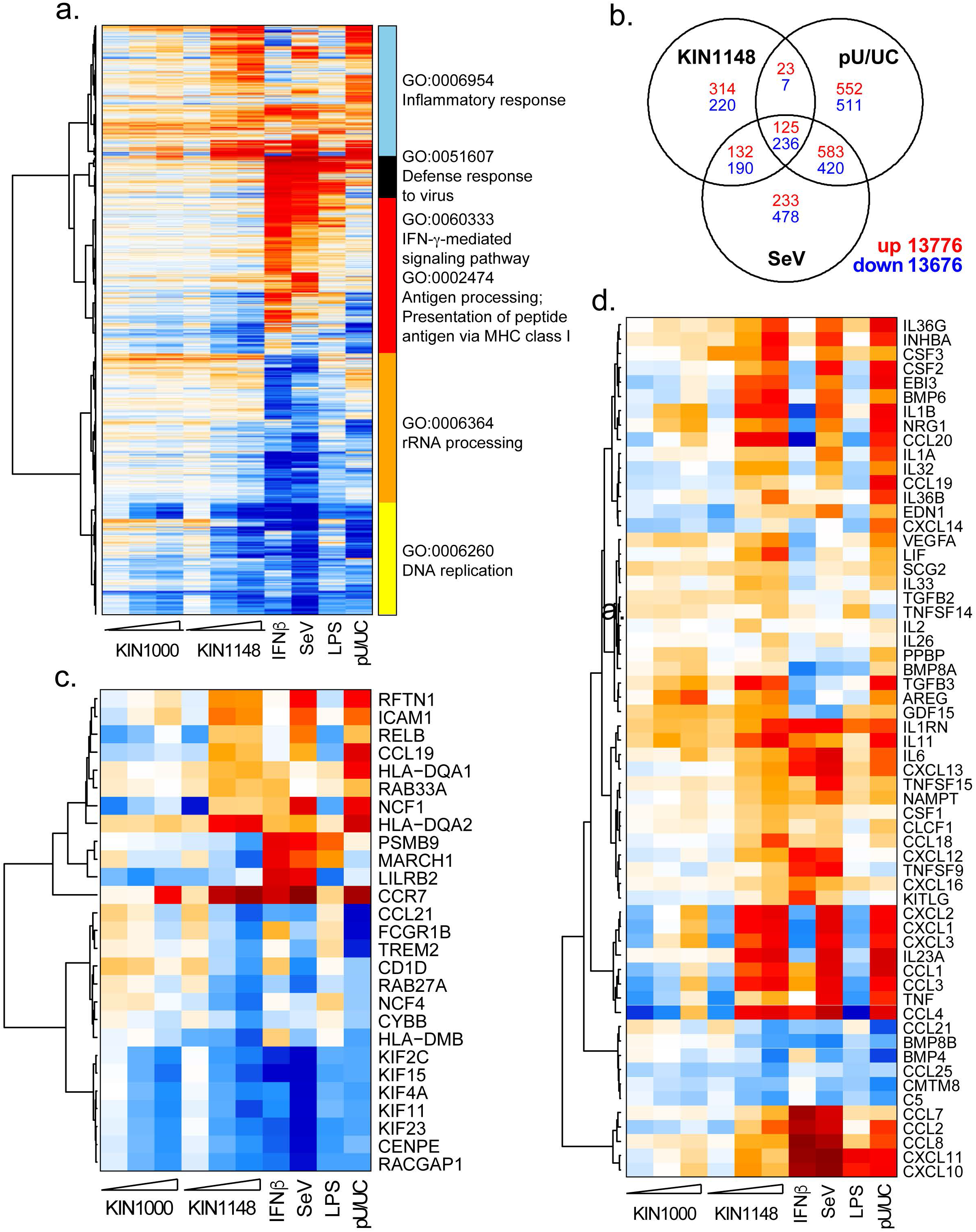
KIN1148 treatment induces a gene expression profile that shares common and unique properties to that achieved with LPS, SeV-infection and polyU/UC RNA transfection. Microarray analysis of THP-1 cells treated with KIN1000 or KIN1148 at concentrations up to 20 μM, 0.5 μg/mL LPS or 100 IU/mL IFNβ as compared to DMSO control. For a RIG-I specific gene expression signature, polyU/UC PAMP RNA-transfected cells were compared to those transfected with xRNA whereas SeV-infected cells were compared to mock-infected cells. Differential gene expression is defined as at least a 2-fold change in expression with a Benjamini-Hochberg corrected p-value <0.01 as compared to the respective negative controls. **(a)** Heatmap showing all genes differentially expressed in at least one treatment with hierarchical clustering and classification by the most highly-enriched gene ontology (GO) biological process. **(b)** Venn diagram showing numbers of differentially expressed genes, up (red) and down (blue), that are shared and unique among cells stimulated with KIN1148, polyU/UC or SeV. (c-d) Analysis showing global gene expression profile with Gene Ontology clustering of enriched genes. Heatmap of genes whose expression is perturbed by compound treatment and designated by gene clustering as **(c)** antigen presentation or **(d)** cytokines.

We plotted all genes differentially expressed in at least one treatment by heatmap and performed hierarchical clustering using spearman correlation as a distance measure (Figure 2a). Patterns of gene expression across doses demonstrate KIN1148 induces changes in gene expression more potently than its parent KIN1000. Consistent with its role as a RIG-I agonist, KIN1148 promoted the expression of genes involved in the inflammatory response, defense response to virus, and a subset of genes involved in antigen processing and presentation. Many genes regulated by KIN1148 are also differentially expressed upon polyU/UC transfection or SeV infection (Figure 2b).

Further inspection of genes whose expression is perturbed by compound treatment demonstrate that KIN1148 induces the expression of many genes designated as “antigen presentation” (Figure 2c) and “cytokines” (Figure 2d) in a dose-dependent manner. Notably, KIN1148-treatment induces a cytokine expression profile that more closely resembles that induced by SeV-infection, polyU/UC transfection or LPS-treatment as opposed to IFNβ-treatment. Although the microarray included assessments of IFNA4, IFNA7, IFNA21, IFNB1, IFNW1 and IFNG, their expression was not detected following KIN1148 treatment (Table S1), while induction of IRF3 target genes were clearly represented. This outcome links KIN1148 binding to RIG-I with downstream IRF3 activation and induction of IRF3-target genes rather than a response attributed merely to induction of type I or type III IFNs. In sum, KIN1148 induces a gene expression profile that is consistent with RIG-I signaling activation in a manner of IRF3 activation that is distinct from IFN-dependent responses, and demonstrates an overall response representing antiviral innate immune activation and adaptive immune priming gene induction.

### KIN1148 augments protection against H5N1 and H1N1 and induces adaptive immune responses following vaccination

We next assessed KIN1148 for the ability to adjuvant an influenza A virus split vaccine (IAV-SV) derived from A/Cal/04/09, the 2009 pandemic H1N1 influenza virus (H1-SV) or PR8/H5N1 (H5-SV), a reassortant PR8 influenza virus containing hemagglutinin and neuraminidase from the highly pathogenic H5N1 avian influenza virus, A/Vietnam/1203/2004. Mice were immunized intramuscularly (i.m.) with a suboptimal dose of H1-SV or H5-SV in combination with PBS, vehicle or KIN1148 and challenged with 5x LD50 (dose required to achieve 50% morbidity) of homologous influenza virus (PR8/H5N1 or A/Cal/04/09) 30 days following immunization. Mice that received IAV-SV in combination with KIN1148 showed significantly decreased mortality, reduced illness scores, and weight loss following lethal PR8/H5N1 (Figure 3a) or lethal A/Cal/04/09 (Figure 3b) challenge compared to IAV-SV + PBS or IAV-SV + vehicle control groups. This enhancement in survival is accompanied by a significant reduction in pulmonary virus titer on day 5 following challenge with A/Cal/04/09 (Figure 3c). These data demonstrate that KIN1148 adjuvants H5-SV and H1-SV to reduce pulmonary virus titer and protection and upon challenge.

**Figure 3.**
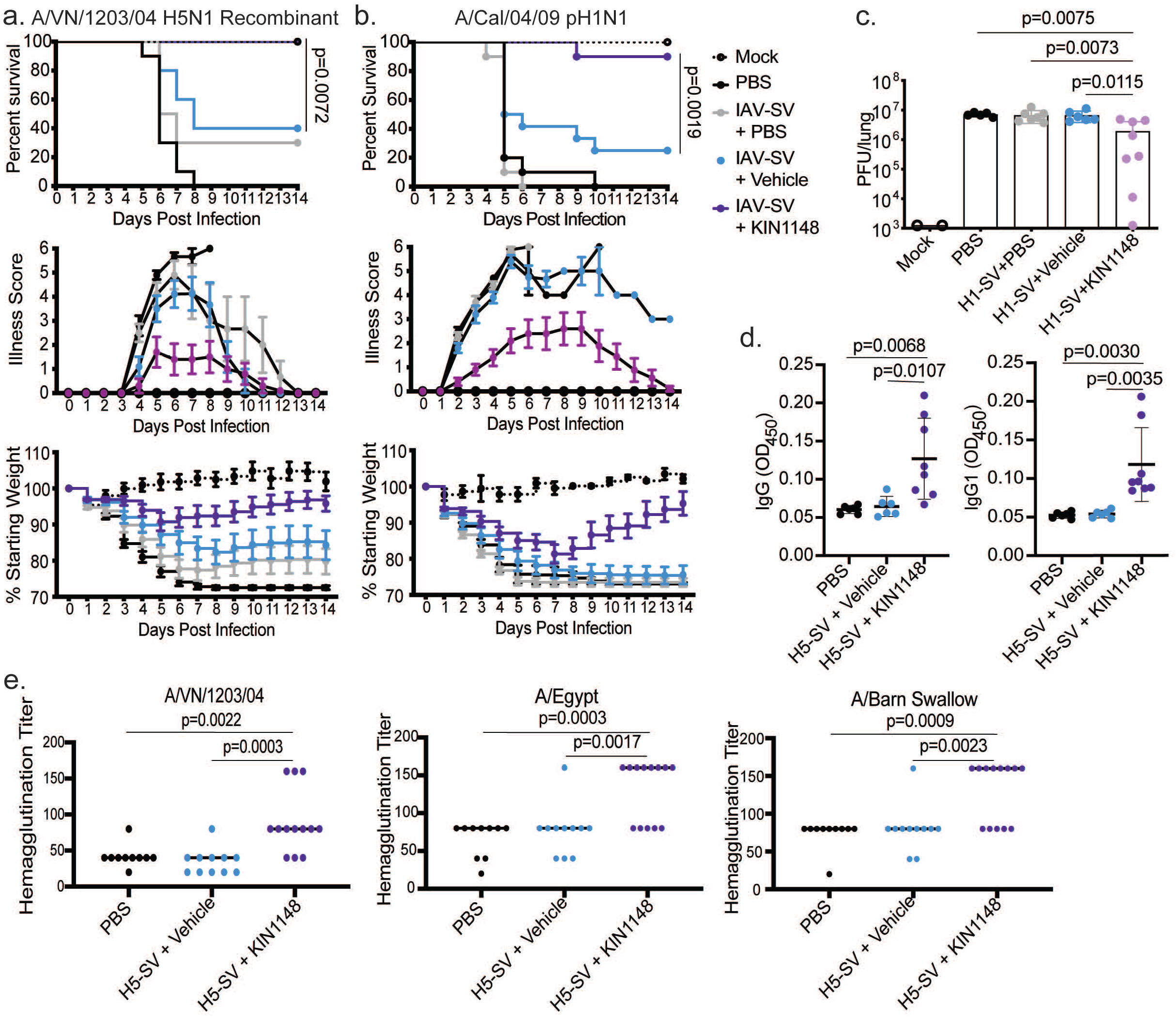
KIN1148 adjuvants influenza virus vaccination induce humoral responses and confers protection against H1N1 and H5N1 infection. **(a-c)** C57BL/6J mice (n=10 mice per group except mock where n=5) were immunized intramuscularly with **(a)** A/VN/1203/04 H5N1 6+2 SV (H5-SV) or **(b)** A/Cal/04/09 H1N1 SV (H1-SV) in combination with PBS (IAV-SV + PBS, grey circles), blank liposome (IAV-SV + Vehicle, blue circles), KIN1148 formulated in liposome (IAV-SV + KIN1148, purple circles) or PBS alone (black circles). On day 30 following vaccination mice were challenged intranasally with 5x LD_50_ of the homologous virus used for vaccination and monitored for survival (top), clinical illness (middle), and weight loss (bottom). **(c)** Mice were immunized and challenged as described in a using H1-SV. On day 5 p.i., lungs were harvested and virus titers were determined by plaque assay. n=2(mock)-8 mice/group with data representative of two independent experiments. **(d)** C57BL/6J mice were immunized intramuscularly PBS alone (PBS, black circles), H5-SV in combination with blank liposome (H5-SV + Vehicle, blue circles), or H5-SV with KIN1148 formulated in liposomes (H5-SV + KIN1148, purple circles). Mice were boosted on day 14 and serum was collected 5 days later for analysis of A/VN/1203/04-specific antibodies. n=6-8 mice/group with data from two pooled experiments shown. **(e)** C57BL/6J mice were immunized as described in **d**. Mice were administered a homologous boost on day 14 and serum was collected on day 19. The ability of antibodies in the serum to prevent hemagglutination of homologous (A/VN/1203/04 H5N1 6+2) or heterologous (A/Egypt/N03072/10 H5 7+1, A/Barn Swallow/HK/D10-1161/2010 7+1) recombinant avian influenza viruses was determined. n=10-13 mice/group with data from two pooled experiments shown.

To determine which protective immune responses were adjuvanted by KIN1148 in the context of IAV-SV to confer protection, we assessed serum antibody responses 5 days following D0/D14 i.m. prime/boost with H5-SV plus PBS, vehicle, or KIN1148. We observed a significant increase in total IgG and IgG1 in the serum of mice immunized with H5-SV + KIN1148 compared to immunization with PBS or vehicle (Figure 3d). KIN1148 also significantly increased hemagglutination inhibition of homologous A/VN/1203/04, but also two other H5N1 recombinant strains, A/Egypt/N03072/10 PR8 recombinant and A/Barn Swallow/HK/D10-1161/2010 PR8 recombinant (Figure 3e). Together, these studies show KIN1148 adjuvants to induce broadly neutralizing antibody responses.

Given the ability of KIN1148 to adjuvant antibody responses in the context of IAV-SV, we assessed whether KIN1148 could augment cell-mediated immunity, potentially enhancing broad protection against IAV. Utilizing the prime/boost strategy described for Figure 4, we observed that KIN1148 significantly increases both the frequency and numbers of germinal center (GC) B cells in the draining popliteal lymph node following vaccination with H5-SV (Figure 4a). KIN1148 also significantly increased IAV-specific CD4^+^ and CD8^+^ T cell responses in the spleen following H5-SV vaccination (Figure 4b and 4c). We observed similar results when A/Cal/04/09 SV was administered with KIN1148 (data not shown). Importantly, these data demonstrate that KIN1148 can augment H5-SV and H1-SV to induce broadly protective cellular immune responses typically not generated following vaccination with IAV-SV alone.

**Figure 4.**
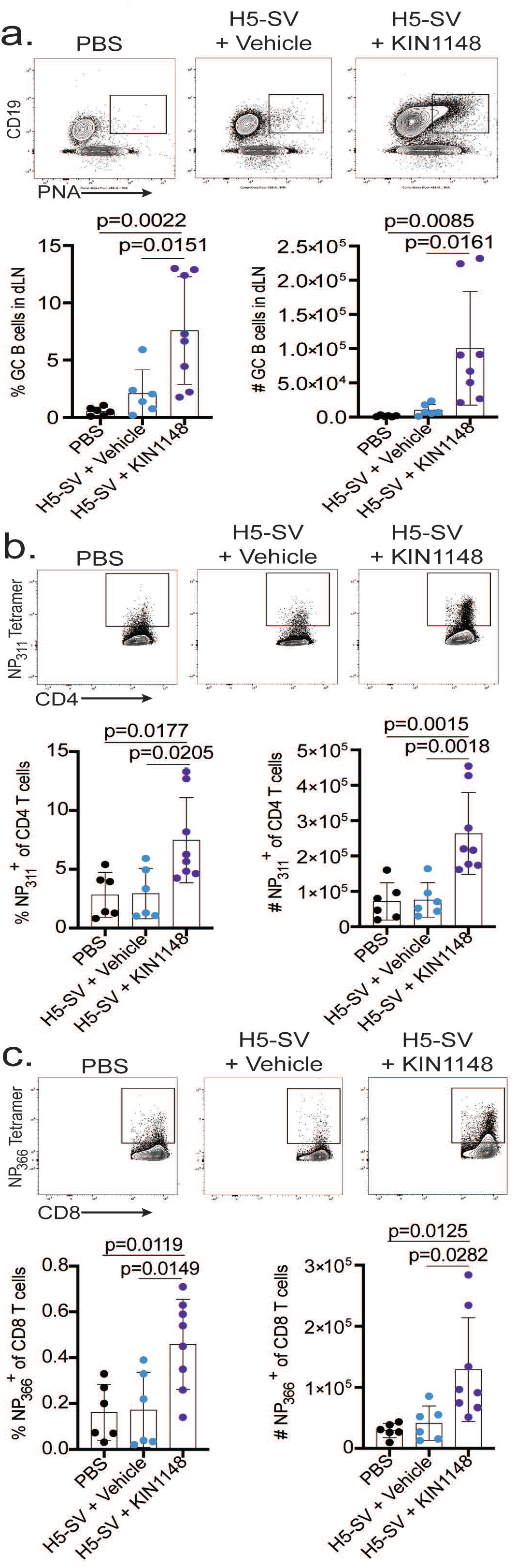
KIN1148 enhances cellular immune responses following H5N1 vaccination. C57BL/6J mice were immunized intramuscularly with PBS (black circles), H5-SV in combination with blank liposome (H5-SV + Vehicle) or H5-SV with KIN1148 formulated in liposomes (H5-SV + KIN1148, purple circles). Mice were administered a homologous boost on day 14 and lung-draining lymph nodes and spleen were collected on day 19. **(a)** The frequency and number of PNA+ GC B cells in dLN was determined by flow cytometry. **(b)** IAV-specific CD4^+^ T cell responses were assessed by flow cytometry. **(c)** IAV-specific CD8^+^ T cell responses were assessed by flow cytometry. n=6-8 mice/group with data from two pooled experiments shown.

### KIN1148 promotes human DC maturation and T cell maturation

To define mechanisms of KIN1148 adjuvant action in human immune cells, we evaluated its effects on DC maturation by treating human monocyte-derived DC (moDC) with DMSO, LPS or KIN1148 for 18 hrs. KIN1148-treated cells exhibit greater cell surface expression of the DC co-stimulatory molecules CD83 and CD86 as compared to DMSO (Figure 5a).

**Figure 5.**
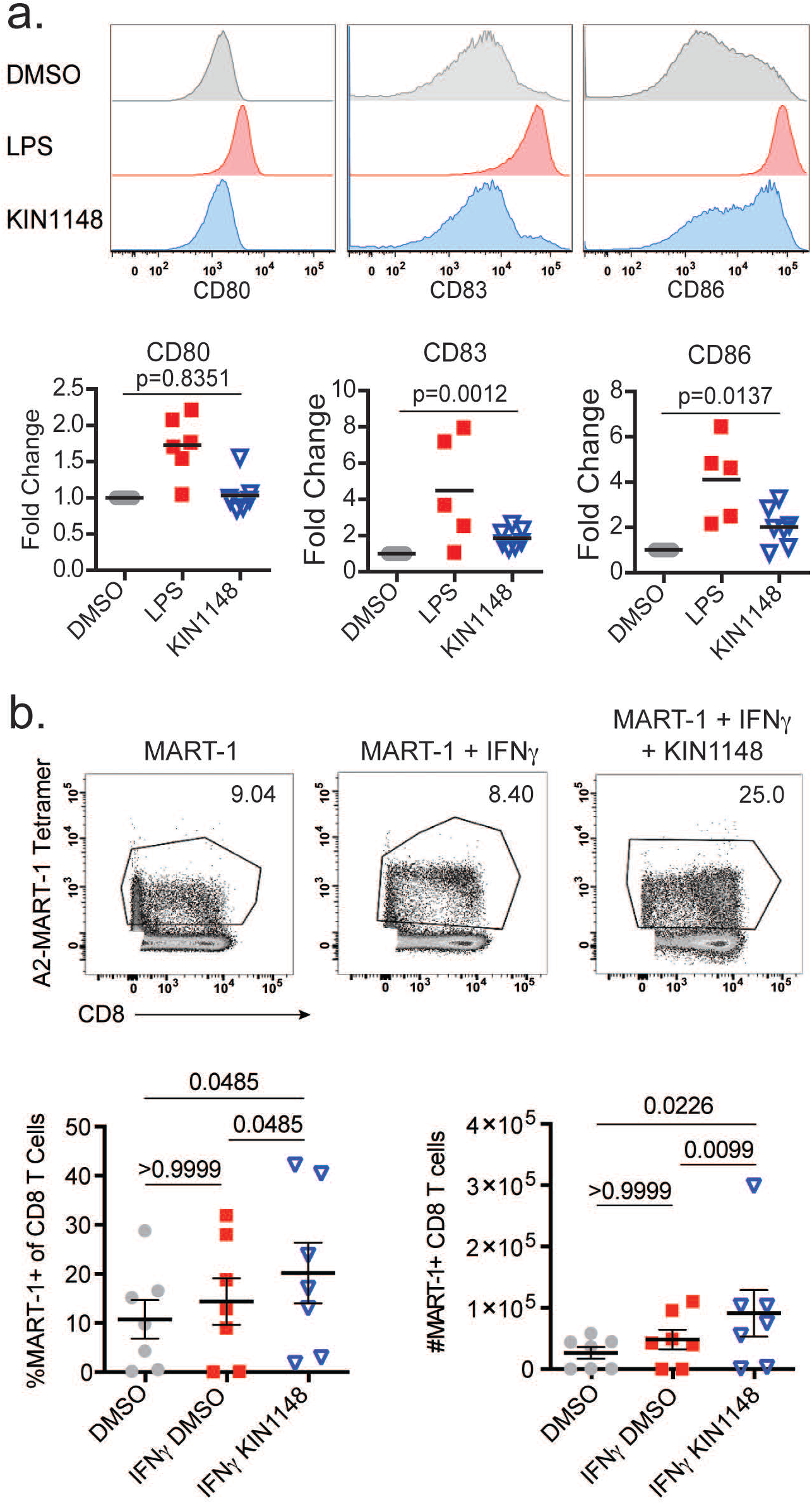
KIN1148 promotes DC maturation and human T cell proliferation. **(a)** Flow cytometry analysis of maturation markers expressed on the cell surface of human monocytic DCs after 18hr treatment with DMSO, LPS or KIN1148. Histograms **(a, top)** of CD80, CD83, and CD86 on human monocytic DCs from a representative experiment. Graphs **(a, bottom)** showing the expression of each maturation marker from at least 6 different healthy donors, expressed as fold change in mean fluorescence intensity (MFI) as compared to DMSO control. **(b)** PBMCs from healthy HLA-A0201^+^ donors were pulsed with Melan-A/MART-1 peptide in combination with DMSO or KIN1148 in the presence of IFNγ for 24 hrs and then removed from the treatments to be co-cultured with autologous T cells in the presence of IL-2. Cells were collected 11 days later and analyzed by flow cytometry for CD8^+^ T cells that stain positive with the corresponding Melan A/MART-1 tetramer. Density plot from a representative experiment showing CD8 and MART-1 tetramer staining and gating strategy (b, top). Graphs (b, bottom) showing the frequency and numbers of MART-1-tetramer^+^ CD8 T cells from 7 donors.

Given the ability of KIN1148 to lead to THP-1 gene responses associated with antigen presentation, induce costimulatory molecule upregulation of moDC, and adjuvant H5-SV to induce IAV-specific CD4^+^ and CD8^+^ T cell responses in our murine model, we evaluated the ability of KIN1148 to enhance human antigen-specific CD8^+^ T cell responses. To simulate conditions similar to prime immunization *ex vivo*, we evaluated CD8^+^ T cell responses to the Melan-A protein using PBMCs from healthy donors. The precursor frequency against this antigen is relatively high in HLA-A0201^+^ individuals and have been shown to represent a normal naive CD8^+^ T cell population in healthy individuals^41–43^, allowing for testing of KIN1148 to augment CD8^+^ T cell activation. PBMCs were pulsed with a Melan-A/MART-1 peptide in combination with DMSO, IFNγ with DMSO or IFNγ with KIN1148 and cultured for 11 days. As anticipated, Melan-A/MART-1-specific CD8^+^ T cell numbers represent a measurable fraction of total CD8^+^ T cells following Melan-A/MART-1 peptide stimulation with or without IFNγ (Figure 5b). This population was significantly increased following co-culture with PBMCs pulsed with Melan-A/MART-1 peptide in combination with KIN1148+IFNγ as compared to DMSO or DMSO+IFNγ controls (Figure 5b). Taken together, our data indicates that KIN1148 promotes antigen presentation leading to greater activation of naive human CD8^+^ T cells.

## Discussion

We report here the identification of KIN1148, a benzobisthiazole small molecule RIG-I agonist that induces IRF3 and NFκB dependent innate immune activation to confer adjuvant activity. KIN1148 was designed through structural-activity relationship (SAR) studies to have improved solubility and pharmacokinetic properties compared to KIN1000, an IRF3-activating small molecule compound identified through screening^23^.

Although it is possible that KIN1148 may direct signaling via additional cellular targets, we provide evidence showing that KIN1148 directly interacts with RIG-I in biochemical assays and cells, and signals for innate immune gene induction in a RIG-I-dependent manner. KIN1148 induces a pattern of gene expression resembling infection with SeV (an RNA virus that activates RIG-I) and by transfection of polyU/UC (a RIG-I-specific RNA agonist). However, we propose that KIN1148 interacts with RIG-I distinctly from PAMP RNA, which binds to the RIG-I RD to induce conformational changes that releases its CARDs to interact with downstream signaling cofactors. Our data demonstrate that KIN1148 binds not only RD, but also the helicase domain of RIG-I. Our studies further reveal that KIN1148 does not compete with ATP or AMP-PNP for binding and does not lead to independent ATP hydrolysis despite signalosome formation and induction of gene responses with cellular treatment of KIN1148 alone. Together, this suggests a potentially novel mechanism for non-canonical RIG-I activation and highlights the potential usefulness of KIN1148 as an adjuvant.

*In vivo*, when administered to mice in conjunction with suboptimal doses of vaccine, KIN1148 was able to enhance the protection of mice from subsequent analogous challenge by a highly pathogenic H5N1 or pandemic H1N1 influenza virus beyond vaccine alone. Previous studies have shown that the addition of RIG-I-activating RNA to vaccines, including pieces of viral RNA and synthetic polyI:C, can similarly enhance the protection of mice above vaccine alone by leading to a Th2-biased responses and enhancement of virus-specific antibody responses^17,24^. Importantly, our results demonstrate that KIN1148 promotes antigen-specific T cell responses *in vivo* in mice and *ex vivo* in humans. As administration of IAV-SV alone does not typically induce T cell-mediated immunity, our results suggest that KIN1148 not only adjuvants vaccine-mediated protection but may do so in a way that broadens immunity against heterosubtypic or drifted IAV strains. Mechanistically, our *ex vivo* studies further demonstrate that KIN1148 promotes DC maturation and their secretion of cytokines that functionally leads to the recruitment of a variety of immune cells.

Overall, this study reveals a small molecule compound which directly binds to RIG-I, inducing a non-canonical response that features IRF3 activation in the absence of type I and III IFN expression, leading to DC activation and enhanced antigen-specific T cell activation of human primary cells. KIN1148 in a liposomal formulation not only adjuvants protection in a murine model of H5N1 and pH1N1 infection but leads to the enhancement of broadly cross-protective antibody and T cell responses.

## Methods

### Cells and viruses

The HEK293 and MDCK cell lines were maintained in complete Dulbecco’s Modified Eagle Medium (cDMEM), supplemented with 10% fetal bovine serum (FBS), L-glutamine, sodium pyruvate and nonessential amino acids. The THP-1 (human monocytic) cell line was maintained in complete RPMI (cRPMI), supplemented with 10% heat inactivated FBS, L-glutamine, sodium pyruvate and nonessential amino acids. THP-1 cells were differentiated in cRPMI supplemented with 40 nM phorbol12-myristate13-acetate (PMA) for 30 hrs prior to being used in experiments. Human peripheral blood mononuclear cells (PBMCs) or monocytes were isolated from buffy coat of healthy human donors (Biological Specialty Corp) by density centrifugation in Ficoll®-Paque PLUS (GE Healthcare Life Sciences). Monocytes were enriched either through positive selection using CD14 MicroBeads (Miltenyi Biotech) or by adherence to tissue culture-treated plates. Monocytes were cultured in cRPMI supplemented with recombinant human IL-4 and GM-CSF (Peprotech) to generate immature monocyte-derived dendritic cells (Mo-DCs), or cDMEM supplemented with recombinant human M-CSF (Peprotech) to generate macrophages. Bone marrow cells isolated from the bones of adult C57BL/6J mice were differentiated into bone marrow-derived macrophages (BMM) in cDMEM supplemented with recombinant mouse M-CSF (Peprotech) or into bone marrow-derived dendritic cells (BMDC) in cRPMI supplemented with recombinant mouse IL-4 and GM-CSF (Peprotech). Influenza virus strains A/California/04/2009 (H1N1), A/VN/1203/04 H5N1 recombinant containing HA with an attenuating mutation in the polybasic cleavage site and NA from the H5N1 virus and all other segments from A/Puerto Rico/8/1934 (PR8), and other H5 recombinants (A/Egypt/N03072/10 and A/Barn Swallow/HK/D10-1161/2010) containing HA with an attenuating mutation in the polybasic cleavage site from the H5 virus and all other segments from A/Puerto Rico/8/1934 (PR8) were propagated in eggs and titered in MDCK cells using plaque assay as described previously described^44^. Sendai virus strain Cantell was obtained from Charles River Laboratories.

### Compounds

Compounds were synthesized by Life Chemicals, Inc. and solubilized in 100% DMSO and kept frozen as 10 mM stocks. The compounds were stored in small aliquots to prevent multiple freeze-thaws and were stepwise diluted to reach the desired concentration in 0.5% (v/v) DMSO for all treatments. Liposomal formulation of KIN1148 for *in vivo* use was performed by InImmune, Inc.

### Preparation of vaccine stocks for *in vivo* studies

Influenza virus A/Cal/04/09 and PR8/H5N1 split vaccine stocks were prepared as previously described^45, 46^. Viral protein content was verified by immunoblot and SDS-PAGE with Coomassie-blue staining, using stocks of BSA of known concentration as standard.

### *In vivo* vaccine studies

Adult C57BL/6J mice were purchased from Jackson Laboratories. For IAV challenge studies, mice were immunized once i.m. with H5- or H1-SV in combination with PBS, KIN1148 in a lipid-based liposomal formulation or the liposomal formulation (vehicle) alone. The mice were challenged 30 days later with 5x LD50 homologous IAV strain PR8/H5N1 or A/Cal/04/09, administered via intranasal instillation in both nares. For morbidity studies, mice were monitored daily for changes in body weight and clinical scores. Mice that were moribund, exhibited 30% or greater weight loss after IAV-infection were euthanized. Additional groups of mice were harvested on day 5 post infection to determine pulmonary virus titer by plaque assay. For experiments analyzing immune responses, C57BL/6J mice were immunized intramuscularly with H5-SV in combination with PBS (no adjuvant control), blank liposome (vehicle control), KIN1148 formulated in liposome or PBS alone (no vaccine control). Mice were boosted on day 14 and harvested 5 days later for serum analysis of antibody responses ELISA, H5N1 hemagglutination inhibition, and GC B cell and IAV-specific T cell responses by flow cytometry. All animal handling and experiments were conducted using protocols approved by the University of Washington Institutional Animal Care and Use Committee (IACUC).

### Antibodies

The following primary antibodies were used for immunoblot detection: Rabbit anti-RIG-I (969, raised in rabbit against a RIG-I CARD peptide sequence)^47^, rabbit anti-MDA5 (Enzo Life Sciences), rabbit anti-LGP2 (IBL-America), rabbit anti-DHX15 (Abcam), rabbit anti-TRIM25 (Cell Signaling), rabbit anti-IKKγ (Santa Cruz), rabbit anti-MAVS (Novus Biologicals), rabbit anti-IRF7 (Cell Signaling), rabbit anti-IRF3 (KINETA, Inc.), rabbit anti-IRF3 phospho-Serine 396 (Cell Signaling), rabbit or mouse anti-FLAG (Sigma), mouse anti-TBK1 (Imgenex), mouse anti-GAPDH (Abcam) and mouse anti-actin (EMD Millipore). Horseradish peroxidase (HRP)-conjugated secondary antibodies were obtained from Jackson ImmunoResearch. Alexa Fluor 488- and conjugated secondary antibodies were obtained from Thermo Fisher Scientific. Near-infrared fluorescent dye (IRDye®)-conjugated secondary antibodies were obtained from LI-COR. The following fluorochrome conjugated anti-mouse antibodies were used for flow cytometry analysis: CD4 (BioLegend), CD8 (BioLegend), CD19 (BioLegend), PNA (Vector Labs), CD3 (BioLegend), CD45.2 (BioLegend). The following fluorochrome conjugated anti-human antibodies were purchased from BioLegend and used for flow cytometry analysis: CD11c, CD80, CD83, CD86, HLA-DR, CD3, and CD8. Live cells were discriminated using Fixable Viability Dye eFluor780 (eBioscience). MelanA, NP366, and NP311 tetramers were acquired from the NIH Tetramer core.

### Antibody response quantification

*ELISA to detect influenza-specific antibodies*. ELISA to detect IAV-specific IgG and IgG1 was performed as previously described^24^. Briefly, ELISA plates (Costar) were coated with UV-inactivated recombinant H5N1 virus. Dilutions of mouse serum following prime boost as described in Figure 3c were plated. Anti-mouse IgG and IgG1 conjugated to biotin (Southern Biotech), High Sensitivity Streptavidin-HRP (ThermoFisher), and 1-Step Ultrafast TMB substrate (ThermoFisher) were used to detect antibody isotypes. *Hemagglutination inhibition assay* was performed as previously described^48^. Briefly, dilutions of serum treated with receptor destroying enzyme (Denkin-Seiken) are incubated with 160 HAU/mL of influenza virus. Subsequently, this virus/serum mixture is incubated with 0.5% chicken red blood cells (Rockland) and hemagglutination inhibition was measured after 30 minutes.

### RT-PCR quantitation of innate immune genes

Cells were harvested in RLT buffer (Qiagen), and total cellular RNA was purified using the RNeasy kit (Qiagen). cDNA was synthesized from the purified RNA by both random and oligo(dT) priming using the iScript cDNA synthesis kit (Bio-rad). RNA levels were measured by the SYBR Green relative quantitation method using either a 7300 or Viia 7 RT-PCR machine (Applied Biosystems). Samples were normalized by subtracting the respective CT values of housekeeping genes glyceraldehyde-3-phosphate dehydrogenase (GAPDH) or human acidic ribosomal protein (HuPO), and fold induction of specific genes calculated relative to DMSO treatment control. Primer sequences are available upon request.

### Biotin-compound pull-down and immunoblot analyses

Cells were trypsinized and collected in RIPA buffer (50 mM Tris-Cl pH 7.5, 150 mM NaCl, 5mM EDTA, 1% NP-40, 0.5% sodium deoxycholate, 0.1% SDS) supplemented with a cocktail of protease inhibitors (Sigma) and phosphatase inhibitors (Calbiochem) and okadaic acid (Calbiochem). Following cell lysis, nuclear material was removed by centrifugation at 15,000 x g for 10 minutes at 4°C. Cell lysates were quantified by BCA (Thermo Fisher Scientific), analyzed on a denaturing Tris-HCl polyacrylamide gel, and transferred onto nitrocellulose membranes. Cellular proteins of interest were detected by immunoblot analysis using specific primary antibodies described earlier and secondary antibodies conjugated to either HRP or near-infrared fluorescent dyes (IRDye®; LI-COR).

Immunoblots developed with HRP-conjugated secondary antibodies were visualized by chemiluminescence on X-ray film and quantitated using the ImageJ software [see the National Institutes of Health ImageJ website http://imagej.nih.gov/ij/ and ^49^].

Immunoblots developed with IRDye®-conjugated secondary antibodies were captured and quantitated using the Odyssey CLx imaging system with Image Studio™ 5.x Software.

### ATPase Assay

ATPase Assays were performed as previously described^23^. Briefly, 2 pmol RNA (xRNA or pU/UC) was mixed with 5 pmol recombinant purified RIG-I and increasing concentrations of recombinant purified KIN1148 (0 pmol, 1.25 pmol, 2.5 pmol, 5 pmol, 10 pmol). 1 mM ATP (Sigma) was added and incubated for 15 min at 37 degrees C. ATPase reaction buffer and BIOMOL Green (Enzo Life Sciences) was added to each sample and read at OD_630_ using a microplate reader.

### Microarray analysis

Differentiated THP-1 cells were treated with small molecule compounds (KIN1000 and KIN1148 at 0.625, 2.5 or 10 μM), 25 IU/mL IFNβ diluted in cRPMI supplemented with 0.5% (v/v) DMSO, or treated with cRPMI supplemented with 0.5% (v/v) DMSO alone (DMSO). Control cells infected with 25 HAU/mL SeV were maintained in cRPMI supplemented with 0.5% (v/v) DMSO after removal of the virus inoculum. Additional cells were transfected with 2 μg/mL polyU/UC RNA (RIG-I-specific PAMP RNA derived from the hepatitis C virus genome ^8^) or control XRNA (RNA segment derived from the hepatitis C virus genome adjacent to and equivalent in length to polyU/UC that cannot induce RIG-I-dependent innate immune signaling^8^) using the *Trans*IT®-mRNA Transfection kit (Mirus Bio LLC). Cells were collected in RLT buffer (Qiagen) 20 hours after treatment or infection. Total RNA purified using RNeasy kit (Qiagen) was submitted to Labcorp Seattle (previously Covance) for microarray analysis using Agilent SurePrint G3 Human Genome Microarrays (version 2). Array data was processed by the Gale lab using R (version 3.2.1)/Bioconductor (version 3.1) [See website by the R Development Core Team at https://www.r-project.org and reference^50^]. Raw data was quantile normalized followed by linear modeling using the limma package (version 3.24.15)^51^. Differential gene expression was defined as at least a 2-fold change in expression with a Benjamini-Hochberg corrected p-value < 0.01 as compared to the appropriate negative control: xRNA transfection for polyU/UC, mock-infected cells for SeV and DMSO for all other samples. The gene expression heatmap was clustered using Spearman correlation distances, and Gene Ontology Biological Processes enriched in lists of genes mapping to such clusters were determined using DAVID 6.7^52,53^. Genes with predicted IRF7 binding sites according to the UCSC Genome Browser database were identified using Enrichr^54^. Finally, genes mapping to the Reactome *Homo sapiens* IFNα/β signaling pathway (R-HAS-909733) were identified using InnateDb^55^. The microarray data discussed in this publication have been deposited in the NCBI Gene Expression Omnibus^56^ and is accessible through GEO accession code: GSE205964.

### DC maturation assay

Monocytes were separated from PBMCs of healthy individuals (Biological Specialty Corps) using anti-CD14 magnetic beads. The monocytes were cultured in cRPMI supplemented with recombinant human IL-4 and GM-CSF (Peprotech) to induce DC differentiation. Day 6-7 immature monocyte derived DCs are exposed for 16-18 hrs to cRPMI supplemented with small molecule compound (KIN1000 or KIN1148; at indicated doses) or 0.5 μg/mL LPS in a final concentration of 0.5% (v/v) DMSO, or cRPMI supplemented with a final concentration of 0.5% (v/v) DMSO alone. DCs were analyzed for expression of maturation markers on the cell surface by flow cytometry using an LSR II (BD Biosciences) and FlowJo v10.1 software (University of Washington Cell Analysis Facility).

### Human *ex vivo* adjuvant assay

PBMCs from healthy HLA-A0201^+^ individuals were cultured in Cell Gro DC medium (CellGenix) supplemented for GM-CSF and IL-4 to induce DC differentiation. After 24h, Melan-A-HLA-A2 epitope peptide^57^ in the presence of 40 ng/mL IFNγ in combination with 0.5% (v/v) final concentration DMSO, 10 μM small molecule compound (KIN1000 or KIN1148) or 0.5 μg/mL LPS in a final concentration of 0.5% (v/v) DMSO was added to the culture medium. 24h later and every 3 days, half of the media was replaced by fresh RPMI supplemented with human serum at a final concentration of 8% and IL-2. On day 11, The CD8^+^ T cells frequency and absolute number specific for Melan-A/MART-1 were assessed by flow cytometry within the CD3^+^CD8^+^ T cell population using Melan-A –HLA-A2 tetramer staining.

### Data analyses

Flow cytometry data was collected on a BD LSRII and analyzed with FlowJo v10. Statistical analyses were performed using GraphPad Prism 7 software (La Jolla). Depending on the number of variables and time points in each experiment, statistical analysis of mean differences between groups was performed by either a Student’s t-test or a multiway analysis of variance (ANOVA) followed by a Bonferroni or Friedman *post hoc* analysis. Kaplan-Meier survival analyses were analyzed by the log-rank test. Specific statistical tests, p values and sample size are indicated in Figure Legends.

## Acknowledgements

We thank Jeffrey Posakony (Kineta, Inc.) for synthesizing biotinylated-KIN1148 and Ran Dong (University of Washington) for providing excellent technical support. Special thanks go to Kristin Bedard, Peter Probst, and Shawn Iadonato from Kineta, Inc. for providing KIN1000 and KIN1148 as well as valuable scientific discussions. We thank Drs. Alison M. Kell (University of New Mexico) and Jacob Yount (Ohio State University) for their insightful feedback during preparation of this manuscript. The HLA-A2/MART-1 class I monomers were obtained through the NIH Tetramer Core Facility. This work was supported by the following National Institutes of Health (NIH) grants and contracts: HHSN272200900035C (M.G), HHSN27220130023C (M.G.), HSSN272201400055C (M.G.), U01AI151698 (M.G.), U19AI100625 Project 3 (M.G.), 5U01AI150747 (P.G.T), 75N93019C00051 (A.G.-S.), 75N93019C00046 (A.G.-S.), P01AI097092 (A.G.-S.), and K22AI146141 (E.A.H.). Some of the work was supported by a cooperative agreement (W81XWH-07-2-0067, W81XWH-11-2-0174, W81XWH-18-2-0040) between the Henry M. Jackson Foundation for the Advancement of Military Medicine Inc. and the U.S. Department of Defense. In addition, this work was supported by the St. Jude Center of Excellence for Influenza Research and Surveillance (HHSN272201400006C), and ALSAC to P.G.T and the CRIPT (Center for Research on Influenza Pathogenesis and Transmission), an NIAID funded CEIRR (Center of Excellence for influenza Research and Response, contract number 75N93021C00014) to A.G.-S. E.A.H. was also supported by American Heart Association Postdoctoral Award 17POST33660907. This research was additionally supported by the Cell Analysis Facility Flow Cytometry and Imaging Core and the Center for Innate Immunity and Immune Disease (CIIID) at the University of Washington. The content of this manuscript is solely the responsibility of the authors and does not necessarily represent the official views of any of the institutions mentioned above, the U.S. Department of the Army or the U.S. Department of Defense or the Henry M. Jackson Foundation for the Advancement of Military Medicine.

## Contributions

This project was initially conceived and directed by Y.L. and M.G. R.I. served as project manager for all studies. E.A.H. and Y.L. designed experiments and analyzed data. C.R.W. developed the computational pipelines and conducted the bioinformatics analyses. E.A.H performed *in vivo* vaccination and challenge studies. C.S. performed the human T cell priming experiment with guidance from L.T. Y.L. and M.K. performed remaining studies. P.G.T. generated vaccine stocks and A.G.-S. provided influenza virus strains. The manuscript was written by E.A.H. and Y.L. and edited by R.I. and M.G. All authors read and approved the final manuscript.

## Supplemental Figure Legends

**Figure S1. KIN1148 binds to RIG-I and signals innate immune gene expression in a RIG-I dependent manner. (a)** Western blot detection of MDA5, RIG-I and ISG56 expression in A549 control (CTRL) and CRISPR-directed RIG-I^−/−^ and MDA5^−/−^ cells after 24hr treatment with 100 IU/mL IFNβ. **(b)** qRT-PCR analysis of IRF3 and NFκB target genes in A549 control or CRISPR-directed RIG-I^−/−^ or MDA5^−/−^ cells at 18hrs after KIN1148 treatment. The data shows average fold induction over DMSO control from at least 3 independent experiments. Error bars show standard deviation. **(c)** Western blot detection of endogenous RIG-I, MAVS and PKR as indicated in HEK293 whole cell lysate (input) and amongst those cellular factors that remain associated with biotin-KIN1148 (K) or bead alone (B) under regular pulldown conditions (high) and conditions of reduced stringency (low). **(d)** Western blot detection of endogenous RIG-I and MAVS in the MEF whole cell lysate (input) and pulldown products that associate with either biotin-fluorescein (F) or biotin-KIN1148 (K) beads. **(e)** Western blot detection of purified recombinant Flag-RIG-I protein in the input and pulldown with biotin-KIN1148 beads in the absence (None) and presence of increasing amounts of KIN1148, AMP-PNP or ATP for competition.

## References

1. Brubaker SW, Bonham KS, Zanoni I, Kagan JC. Innate immune pattern recognition: a cell biological perspective. Annu Rev Immunol. 2015;33:257–90. doi: 10.1146/annurev-immunol-032414-112240. PubMed PMID: 25581309; PMCID: PMC5146691.

2. Cui J, Chen Y, Wang HY, Wang RF. Mechanisms and pathways of innate immune activation and regulation in health and cancer. Hum Vaccin Immunother. 2014;10(11):3270–85. doi: 10.4161/21645515.2014.979640. PubMed PMID: 25625930; PMCID: PMC4514086.

3. Kell AM, Gale M, Jr. RIG-I in RNA virus recognition. Virology. 2015;479-480:110–21. doi: 10.1016/j.virol.2015.02.017. PubMed PMID: 25749629; PMCID: PMC4424084.

4. Loo YM, Gale M, Jr. Immune signaling by RIG-I-like receptors. Immunity. 2011;34(5):680–92. doi: 10.1016/j.immuni.2011.05.003. PubMed PMID: 21616437; PMCID: PMC3177755.

5. Chan YK, Gack MU. RIG-I-like receptor regulation in virus infection and immunity. Curr Opin Virol. 2015;12:7–14. doi: 10.1016/j.coviro.2015.01.004. PubMed PMID: 25644461.

6. Kato H, Takeuchi O, Mikamo-Satoh E, Hirai R, Kawai T, Matsushita K, Hiiragi A, Dermody TS, Fujita T, Akira S. Length-dependent recognition of double-stranded ribonucleic acids by retinoic acid-inducible gene-I and melanoma differentiation-associated gene 5. J Exp Med. 2008;205(7):1601–10. doi: 10.1084/jem.20080091. PubMed PMID: 18591409; PMCID: PMC2442638.

7. Kell A, Stoddard M, Li H, Marcotrigiano J, Shaw GM, Gale M. Pathogen-Associated Molecular Pattern Recognition of Hepatitis C Virus Transmitted/Founder Variants by RIG-I Is Dependent on U-Core Length. J Virol. 2015;89(21):11056–68. Epub 20150826. doi: 10.1128/JVI.01964-15. PubMed PMID: 26311867; PMCID: PMC4621103.

8. Saito T, Owen DM, Jiang F, Marcotrigiano J, Gale M, Jr. Innate immunity induced by composition-dependent RIG-I recognition of hepatitis C virus RNA. Nature. 2008;454(7203):523–7. Epub 2008/06/13. doi: nature07106 [pii] 10.1038/nature07106. PubMed PMID: 18548002; PMCID: 2856441.

9. Schnell G, Loo YM, Marcotrigiano J, Gale M. Uridine composition of the poly-U/UC tract of HCV RNA defines non-self recognition by RIG-I. PLoS Pathog. 2012;8(8):e1002839. Epub 20120802. doi: 10.1371/journal.ppat.1002839. PubMed PMID: 22912574; PMCID: PMC3410852.

10. Coffman RL, Sher A, Seder RA. Vaccine adjuvants: putting innate immunity to work. Immunity. 2010;33(4):492–503. doi: 10.1016/j.immuni.2010.10.002. PubMed PMID: 21029960; PMCID: PMC3420356.

11. He S, Mao X, Sun H, Shirakawa T, Zhang H, Wang X. Potential therapeutic targets in the process of nucleic acid recognition: opportunities and challenges. Trends Pharmacol Sci. 2015;36(1):51–64. doi: 10.1016/j.tips.2014.10.013. PubMed PMID: 25479797.

12. Moyer TJ, Zmolek AC, Irvine DJ. Beyond antigens and adjuvants: formulating future vaccines. J Clin Invest. 2016;126(3):799–808. doi: 10.1172/JCI81083. PubMed PMID: 26928033; PMCID: PMC4767337.

13. Lee S, Nguyen MT. Recent advances of vaccine adjuvants for infectious diseases. Immune Netw. 2015;15(2):51–7. doi: 10.4110/in.2015.15.2.51. PubMed PMID: 25922593; PMCID: PMC4411509.

14. Pulendran B, S Arunachalam P, O'Hagan DT. Emerging concepts in the science of vaccine adjuvants. Nat Rev Drug Discov. 2021;20(6):454–75. Epub 20210406. doi: 10.1038/s41573-021-00163-y. PubMed PMID: 33824489; PMCID: PMC8023785.

15. Reed SG, Tomai M, Gale MJ. New horizons in adjuvants for vaccine development. Curr Opin Immunol. 2020;65:97–101. Epub 20201008. doi: 10.1016/j.coi.2020.08.008. PubMed PMID: 33038865; PMCID: PMC7542129.

16. Ziegler A, Soldner C, Lienenklaus S, Spanier J, Trittel S, Riese P, Kramps T, Weiss S, Heidenreich R, Jasny E, Guzman CA, Kallen KJ, Fotin-Mleczek M, Kalinke U. A New RNA-Based Adjuvant Enhances Virus-Specific Vaccine Responses by Locally Triggering TLR- and RLH-Dependent Effects. J Immunol. 2017. doi: 10.4049/jimmunol.1601129. PubMed PMID: 28077601.

17. Beljanski V, Chiang C, Kirchenbaum GA, Olagnier D, Bloom CE, Wong T, Haddad EK, Trautmann L, Ross TM, Hiscott J. Enhanced Influenza Virus-Like Particle Vaccination with a Structurally Optimized RIG-I Agonist as Adjuvant. J Virol. 2015;89(20):10612–24. doi: 10.1128/JVI.01526-15. PubMed PMID: 26269188; PMCID: PMC4580177.

18. Hochheiser K, Klein M, Gottschalk C, Hoss F, Scheu S, Coch C, Hartmann G, Kurts C. Cutting Edge: The RIG-I Ligand 3pRNA Potently Improves CTL Cross-Priming and Facilitates Antiviral Vaccination. J Immunol. 2016;196(6):2439–43. doi: 10.4049/jimmunol.1501958. PubMed PMID: 26819202.

19. Kulkarni RR, Rasheed MA, Bhaumik SK, Ranjan P, Cao W, Davis C, Marisetti K, Thomas S, Gangappa S, Sambhara S, Murali-Krishna K. Activation of the RIG-I pathway during influenza vaccination enhances the germinal center reaction, promotes T follicular helper cell induction, and provides a dose-sparing effect and protective immunity. J Virol. 2014;88(24):13990–4001. doi: 10.1128/JVI.02273-14. PubMed PMID: 25253340; PMCID: PMC4249139.

20. Mercado-Lopez X, Cotter CR, Kim WK, Sun Y, Munoz L, Tapia K, Lopez CB. Highly immunostimulatory RNA derived from a Sendai virus defective viral genome. Vaccine. 2013;31(48):5713–21. doi: 10.1016/j.vaccine.2013.09.040. PubMed PMID: 24099876; PMCID: PMC4406099.

21. Linehan MM, Dickey TH, Molinari ES, Fitzgerald ME, Potapova O, Iwasaki A, Pyle AM. A minimal RNA ligand for potent RIG-I activation in living mice. Sci Adv. 2018;4(2):e1701854. Epub 20180221. doi: 10.1126/sciadv.1701854. PubMed PMID: 29492454; PMCID: PMC5821489.

22. Bedard KM, Wang ML, Proll SC, Loo YM, Katze MG, Gale M, Jr., Iadonato SP. Isoflavone agonists of IRF-3 dependent signaling have antiviral activity against RNA viruses. J Virol. 2012;86(13):7334–44. doi: 10.1128/JVI.06867-11. PubMed PMID: 22532686; PMCID: PMC3416323.

23. Pattabhi S, Wilkins CR, Dong R, Knoll ML, Posakony J, Kaiser S, Mire CE, Wang ML, Ireton RC, Geisbert TW, Bedard KM, Iadonato SP, Loo YM, Gale M, Jr. Targeting Innate Immunity for Antiviral Therapy through Small Molecule Agonists of the RLR Pathway. J Virol. 2015;90(5):2372–87. doi: 10.1128/JVI.02202-15. PubMed PMID: 26676770; PMCID: PMC4810700.

24. Probst P, Grigg JB, Wang M, Munoz E, Loo YM, Ireton RC, Gale M, Iadonato SP, Bedard KM. A small-molecule IRF3 agonist functions as an influenza vaccine adjuvant by modulating the antiviral immune response. Vaccine. 2017;35(15):1964–71. doi: 10.1016/j.vaccine.2017.01.053. PubMed PMID: WOS:000399850800017.

25. Jing YY, Han ZP, Sun K, Zhang SS, Hou J, Liu Y, Li R, Gao L, Zhao X, Zhao QD, Wu MC, Wei LX. Toll-like receptor 4 signaling promotes epithelial-mesenchymal transition in human hepatocellular carcinoma induced by lipopolysaccharide. BMC Med. 2012;10:98. doi: 10.1186/1741-7015-10-98. PubMed PMID: 22938142; PMCID: PMC3482562.

26. Ding Q, Cao X, Lu J, Huang B, Liu YJ, Kato N, Shu HB, Zhong J. Hepatitis C virus NS4B blocks the interaction of STING and TBK1 to evade host innate immunity. J Hepatol. 2013;59(1):52–8. doi: 10.1016/j.jhep.2013.03.019. PubMed PMID: 23542348.

27. Sun W, Li Y, Chen L, Chen H, You F, Zhou X, Zhou Y, Zhai Z, Chen D, Jiang Z. ERIS, an endoplasmic reticulum IFN stimulator, activates innate immune signaling through dimerization. Proc Natl Acad Sci U S A. 2009;106(21):8653–8. doi: 10.1073/pnas.0900850106. PubMed PMID: 19433799; PMCID: PMC2689030.

28. Zhong B, Yang Y, Li S, Wang YY, Li Y, Diao F, Lei C, He X, Zhang L, Tien P, Shu HB. The adaptor protein MITA links virus-sensing receptors to IRF3 transcription factor activation. Immunity. 2008;29(4):538–50. doi: 10.1016/j.immuni.2008.09.003. PubMed PMID: 18818105.

29. Li K, Chen Z, Kato N, Gale M, Jr., Lemon SM. Distinct poly(I-C) and virus-activated signaling pathways leading to interferon-beta production in hepatocytes. J Biol Chem. 2005;280(17):16739–47. doi: 10.1074/jbc.M414139200. PubMed PMID: 15737993.

30. Chow KT, Gale M, Jr., Loo YM. RIG-I and Other RNA Sensors in Antiviral Immunity. Annu Rev Immunol. 2018;36:667–94. doi: 10.1146/annurev-immunol-042617-053309. PubMed PMID: 29677479.

31. Ng CS, Kato H, Fujita T. Recognition of viruses in the cytoplasm by RLRs and other helicases--how conformational changes, mitochondrial dynamics and ubiquitination control innate immune responses. Int Immunol. 2012;24(12):739–49. doi: 10.1093/intimm/dxs099. PubMed PMID: 23087188.

32. Chiang C, Gack MU. Post-translational Control of Intracellular Pathogen Sensing Pathways. Trends Immunol. 2017;38(1):39–52. doi: 10.1016/j.it.2016.10.008. PubMed PMID: 27863906; PMCID: PMC5580928.

33. Cadena C, Ahmad S, Xavier A, Willemsen J, Park S, Park JW, Oh SW, Fujita T, Hou F, Binder M, Hur S. Ubiquitin-Dependent and -Independent Roles of E3 Ligase RIPLET in Innate Immunity. Cell. 2019;177(5):1187–200.e16. Epub 20190418. doi: 10.1016/j.cell.2019.03.017. PubMed PMID: 31006531; PMCID: PMC6525047.

34. Lu H, Lu N, Weng L, Yuan B, Liu YJ, Zhang Z. DHX15 senses double-stranded RNA in myeloid dendritic cells. J Immunol. 2014;193(3):1364–72. doi: 10.4049/jimmunol.1303322. PubMed PMID: 24990078; PMCID: PMC4108507.

35. Mosallanejad K, Sekine Y, Ishikura-Kinoshita S, Kumagai K, Nagano T, Matsuzawa A, Takeda K, Naguro I, Ichijo H. The DEAH-box RNA helicase DHX15 activates NF-kappaB and MAPK signaling downstream of MAVS during antiviral responses. Science signaling. 2014;7(323):ra40. doi: 10.1126/scisignal.2004841. PubMed PMID: 24782566.

36. Pattabhi S, Knoll M, Gale MJ, Loo YM. DHX15 is a pathogen-recognition receptor that promotes antiviral defense against RNA virus infection. Journal Interferon Cytokine Research2019.

37. Wang P, Zhu S, Yang L, Cui S, Pan W, Jackson R, Zheng Y, Rongvaux A, Sun Q, Yang G, Gao S, Lin R, You F, Flavell R, Fikrig E. Nlrp6 regulates intestinal antiviral innate immunity. Science. 2015;350(6262):826–30. doi: 10.1126/science.aab3145. PubMed PMID: 26494172; PMCID: PMC4927078.

38. Fullam A, Schroder M. DExD/H-box RNA helicases as mediators of anti-viral innate immunity and essential host factors for viral replication. Biochim Biophys Acta. 2013;1829(8):854–65. Epub 2013/04/10. doi: S1874-9399(13)00057-6 [pii] 10.1016/j.bbagrm.2013.03.012. PubMed PMID: 23567047.

39. Garbelli A, Beermann S, Di Cicco G, Dietrich U, Maga G. A motif unique to the human DEAD-box protein DDX3 is important for nucleic acid binding, ATP hydrolysis, RNA/DNA unwinding and HIV-1 replication. PLoS One. 2011;6(5):e19810. doi: 10.1371/journal.pone.0019810. PubMed PMID: 21589879; PMCID: PMC3093405.

40. Gu L, Fullam A, Brennan R, Schroder M. Human DEAD box helicase 3 couples IkappaB kinase epsilon to interferon regulatory factor 3 activation. Mol Cell Biol. 2013;33(10):2004–15. doi: 10.1128/MCB.01603-12. PubMed PMID: 23478265; PMCID: 3647972.

41. Alanio C, Lemaitre F, Law HK, Hasan M, Albert ML. Enumeration of human antigen-specific naive CD8+ T cells reveals conserved precursor frequencies. Blood. 2010;115(18):3718–25. Epub 20100303. doi: 10.1182/blood-2009-10-251124. PubMed PMID: 20200354.

42. Pittet MJ, Valmori D, Dunbar PR, Speiser DE, Liénard D, Lejeune F, Fleischhauer K, Cerundolo V, Cerottini JC, Romero P. High frequencies of naive Melan-A/MART-1-specific CD8(+) T cells in a large proportion of human histocompatibility leukocyte antigen (HLA)-A2 individuals. J Exp Med. 1999;190(5):705–15. doi: 10.1084/jem.190.5.705. PubMed PMID: 10477554; PMCID: PMC2195613.

43. Wölfl M, Greenberg PD. Antigen-specific activation and cytokine-facilitated expansion of naive, human CD8+ T cells. Nat Protoc. 2014;9(4):950–66. Epub 20140327. doi: 10.1038/nprot.2014.064. PubMed PMID: 24675735; PMCID: PMC4312138.

44. Szretter KJ, Balish AL, Katz JM. Influenza: propagation, quantification, and storage. Curr Protoc Microbiol. 2006;Chapter 15:Unit 15G.1. doi: 10.1002/0471729256.mc15g01s3. PubMed PMID: 18770580.

45. Wong SS, Webby RJ. Traditional and new influenza vaccines. Clin Microbiol Rev. 2013;26(3):476–92. doi: 10.1128/CMR.00097-12. PubMed PMID: 23824369; PMCID: PMC3719499.

46. Krammer F, Palese P. Advances in the development of influenza virus vaccines. Nat Rev Drug Discov. 2015;14(3):167–82. doi: 10.1038/nrd4529. PubMed PMID: 25722244.

47. Liu HM, Loo YM, Horner SM, Zornetzer GA, Katze MG, Gale M, Jr. The mitochondrial targeting chaperone 14-3-3epsilon regulates a RIG-I translocon that mediates membrane association and innate antiviral immunity. Cell Host Microbe. 2012;11(5):528–37. doi: 10.1016/j.chom.2012.04.006. PubMed PMID: 22607805; PMCID: PMC3358705.

48. Kaufmann L, Syedbasha M, Vogt D, Hollenstein Y, Hartmann J, Linnik JE, Egli A. An Optimized Hemagglutination Inhibition (HI) Assay to Quantify Influenza-specific Antibody Titers. J Vis Exp. 2017(130). Epub 20171201. doi: 10.3791/55833. PubMed PMID: 29286466; PMCID: PMC5755515.

49. Schneider CA, Rasband WS, Eliceiri KW. NIH Image to ImageJ: 25 years of image analysis. Nature methods. 2012;9(7):671–5. PubMed PMID: 22930834.

50. Gentleman RC, Carey VJ, Bates DM, Bolstad B, Dettling M, Dudoit S, Ellis B, Gautier L, Ge Y, Gentry J, Hornik K, Hothorn T, Huber W, Iacus S, Irizarry R, Leisch F, Li C, Maechler M, Rossini AJ, Sawitzki G, Smith C, Smyth G, Tierney L, Yang JY, Zhang J. Bioconductor: open software development for computational biology and bioinformatics. Genome biology. 2004;5(10):R80. doi: 10.1186/gb-2004-5-10-r80. PubMed PMID: 15461798; PMCID: PMC545600.

51. Smyth GK. Linear models and empirical bayes methods for assessing differential expression in microarray experiments. Stat Appl Genet Mol Biol. 2004;3:Article3. doi: 10.2202/1544-6115.1027. PubMed PMID: 16646809.

52. Huang DW, Sherman BT, Tan Q, Collins JR, Alvord WG, Roayaei J, Stephens R, Baseler MW, Lane HC, Lempicki RA. The DAVID Gene Functional Classification Tool: a novel biological module-centric algorithm to functionally analyze large gene lists. Genome biology. 2007;8(9):R183. doi: 10.1186/gb-2007-8-9-r183. PubMed PMID: 17784955; PMCID: PMC2375021.

53. Huang DW, Sherman BT, Tan Q, Kir J, Liu D, Bryant D, Guo Y, Stephens R, Baseler MW, Lane HC, Lempicki RA. DAVID Bioinformatics Resources: expanded annotation database and novel algorithms to better extract biology from large gene lists. Nucleic Acids Res. 2007;35(Web Server issue):W169–75. doi: 10.1093/nar/gkm415. PubMed PMID: 17576678; PMCID: PMC1933169.

54. Chen EY, Tan CM, Kou Y, Duan Q, Wang Z, Meirelles GV, Clark NR, Ma'ayan A. Enrichr: interactive and collaborative HTML5 gene list enrichment analysis tool. BMC Bioinformatics. 2013;14:128. doi: 10.1186/1471-2105-14-128. PubMed PMID: 23586463; PMCID: PMC3637064.

55. Breuer K, Foroushani AK, Laird MR, Chen C, Sribnaia A, Lo R, Winsor GL, Hancock RE, Brinkman FS, Lynn DJ. InnateDB: systems biology of innate immunity and beyond--recent updates and continuing curation. Nucleic Acids Res. 2013;41(Database issue):D1228–33. doi: 10.1093/nar/gks1147. PubMed PMID: 23180781; PMCID: PMC3531080.

56. Edgar R, Domrachev M, Lash AE. Gene Expression Omnibus: NCBI gene expression and hybridization array data repository. Nucleic Acids Res. 2002;30(1):207–10. PubMed PMID: 11752295; PMCID: PMC99122.

57. Herr W, Schneider J, Lohse AW, Meyer zum Buschenfelde KH, Wolfel T. Detection and quantification of blood-derived CD8+ T lymphocytes secreting tumor necrosis factor alpha in response to HLA-A2.1-binding melanoma and viral peptide antigens. J Immunol Methods. 1996;191(2):131–42. PubMed PMID: 8666832.

