## Supplemental Figure 1 and Supplemental Table 1 for "A small molecule RIG-I agonist serves to adjuvant broad multifaceted influenza virus vaccine immunity"

### Figure S1

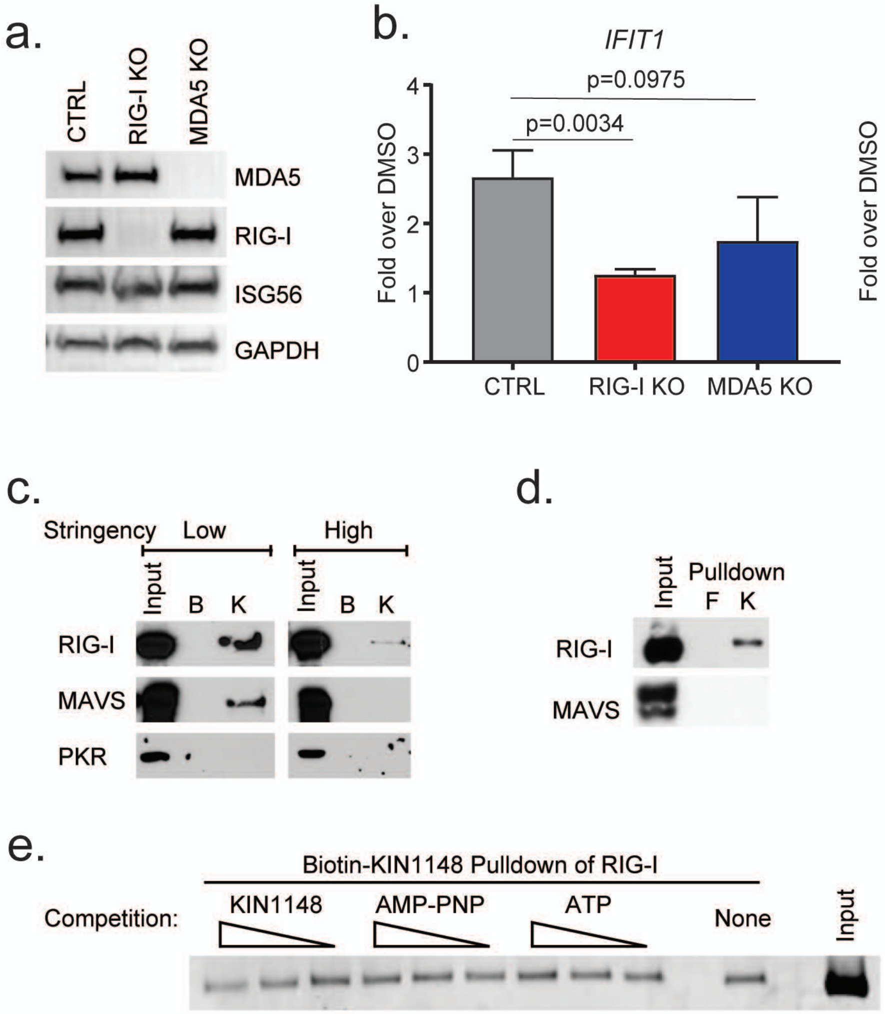
a.
CTRL
RIG-I KO
MDA5 KO
MDA5
RIG-I
ISG56
GAPDH
b.
IFIT1
p=0.0975
p=0.0034
Fold over DMSO
0
1
2
3
4
CTRL
RIG-I KO
MDA5 KO
IL6
p=0.5145
p=0.0002
Fold over DMSO
0
5
10
15
CTRL
RIG-I KO
MDA5 KO
c.
Stringency
Low
High
Input
B
K
Input
B
K
RIG-I
MAVS
PKR
d.
Input
Pulldown
F
K
RIG-I
MAVS
e.
Biotin-KIN1148 Pulldown of RIG-I
Competition:
KIN1148
AMP-PNP
ATP
None
Input

| **Table S1. Expression of IFN and IFN-related genes following innate immune stimulation.**[1] | | | | | | |
|---|---|---|---|---|---|---|
| | KIN1000 | KIN1148 | IFNβ | SeV | LPS | pU/UC |
| IFNA4 | 0 | 0 | 0 | 0 | 0 | +1 |
| IFNA7 | 0 | 0 | 0 | 0 | 0 | +1 |
| IFNA21 | 0 | 0 | 0 | 0 | 0 | 0 |
| IFNB1 | 0 | 0 | 0 | +1 | 0 | +1 |
| IFNW1 | 0 | 0 | 0 | 0 | 0 | +1 |
| IFNG | 0 | 0 | 0 | 0 | 0 | 0 |
| IFNAR1 | 0 | 0 | 0 | 0 | 0 | 0 |
| IFNAR2 | 0 | 0 | 0 | 0 | 0 | 0 |
| IFNGR1 | 0 | -1 | 0 | -1 | -1 | 0 |
| IFNGR2 | 0 | 0 | 0 | 0 | 0 | 0 |

[1]Table S1 shows relative expression levels of various IFN genes and receptors as identified by gene expression following treatment of THP-1 cells with KIN1000, KIN1148, IFNβ, SeV, LPS, or PU/UC as described in Figure 2. (+1) for increased expression, (0) for no difference and (-1) for reduced expression as compared to their respective controls.
